# Acute Administration of Oxytocin in the Functional Recovery of Neurocognitive and Social Deficits Following Juvenile Frontal Traumatic Brain Injury

**DOI:** 10.1101/2025.04.30.651517

**Authors:** Sophia Shonka, Claudia Ford, Michael J. Hylin

## Abstract

**Introduction:** Juvenile traumatic brain injury (jTBI) is one of the leading causes of death and disability in children. The prefrontal cortex (PFC) is most susceptible to injury which leads to deficits in executive function, social behaviors, and cognitive flexibility. Prior research has shown a significant role of the oxytocin (OXT) system in the modulation of social behaviors, and that intranasal OXT (IN-OXT) is potentially neuroprotective. Therefore, we believe IN-OXT could improve functional recovery from a PFC injury.

**Methods:** Animals received a single midline cortical contusion bilaterally damaging the medial PFC (mPFC) and immediately given a single dose of IN-OXT, placebo, or no treatment. Animals were assessed using behavioral and histological measures.

**Results:** The results indicated that jTBI significantly impacted the normal development of the OXT system, likely due to deficits in OXT synthesis in the SON. While IN-OXT alleviated these deficits, it had no impact on neuroinflammation. Similarly, behavioral effects of IN-OXT were not consistent. Mild improvements were observed in spatial learning but there were no improvements in spatial memory or social dominance.

**Discussion:** These results show that IN-OXT increases OXT levels in the brain via pathways originating in the SON and improved spatial learning.

## Introduction

Traumatic brain injury (TBI) is a major global health concern (1,2), with an estimated 64-74 million new cases annually (3). Young children are particularly susceptible to sustaining TBI, most commonly due to falls, with 837,000 new pediatric cases in the United States annually (4,5). The frontal lobe is a frequently damaged region in TBI due to its protuberance, resulting in executive dysfunctions (6,7). These deficits impact all areas of an individual’s life by making participation in work and society challenging, often leading to loss of autonomy (8). Furthermore, TBI results in higher rates of disability in children, with 62% experiencing disability and requiring specialized medical and educational services (4).

Juvenile TBIs (jTBIs) are especially significant because the rehabilitative course overlaps with the normal developmental processes (4,9). Subsequently, children tend to ‘grow into the lesion’, whereby deficits are not fully realized until the damaged brain region finishes developing (10,11). This is especially relevant for the frontal lobe as it has the longest period of development, with anatomical changes measures well into young adulthood (12–14). Subsequently, juvenile TBI results in lower rates of postsecondary education enrollment, independent living, and employment with increased rats of psychiatric disorders and interpersonal difficulties (4,15–18).

Despite the severity of TBI’s impact, there are windows during development where injury response is more favorable (11,19–22). Clinically, older children and teenagers tend to have better global outcomes after injury compared to adults (19). Preclinically, the worst outcomes are observed if injured during the first week of life (20), but there is significant sparing of function if the injury occurs in the second week of life (20,21). Thereafter, rodent outcomes mimic those observed clinically. Thus, there are better ages at which to sustain an injury which provides an avenue through which treatments would likely improve recovery of function.

Oxytocin (OXT) is one such treatment avenue as it has demonstrated neuroprotective effects including social neuroprotection and anti-inflammatory effects (23–25). Thus, the OXT system is a potential route to reverse the secondary injury cascade and enhance recovery following TBI. Indeed, intranasal OXT (IN-OXT) has been shown to be therapeutic in several injury models, including jTBI, stroke, and perinatal brain injury. In jTBI, IN-OXT reduced social novelty deficits following injury by increasing inhibition in the medial prefrontal cortex (mPFC) (25). However, since this model injured the parietal lobe, it is unknown if IN-OXT could rescue or prevent deficits such as this if the frontal lobe is directly damaged by TBI. Meanwhile, stroke models found that IN-OXT immediately following stroke reduced infarct size, decreased neurodegeneration, and mitigated morphological changes (23,26–28). Possibly due to its anti-inflammatory roles, including attenuation of the microglia response (26,28,29) and reducing pro-inflammatory gene expressions (28). Finally, IN-OXT mitigated free-radical production and protected against oxygen-glucose deprivation in perinatal injury models (26,30). Given the significant inflammatory response following TBI, IN-OXT may be a promising treatment against the pathophysiological and behavioral effects of TBI in children. However, very little research has been done on the effects of TBI on the development of the OXT system nor the potential efficacy of IN-OXT as a treatment for inflammation following TBI. Furthermore, no research has been done on how reducing or preventing inflammation using IN-OXT impacts neurocognitive and social deficits that are commonly observed following injury. Thus, this study was conducted to investigate the impact of jTBI on the OXT system and inflammation in rats during the acute phase following injury and evaluate the potential efficacy of IN-OXT as a treatment. To study this, we first examined the impact of jTBI on the normal development of the OXT system by examining neuropathology at different developmental time points. Then, we administered IN-OXT to a second cohort of animals to assess chronic neurocognitive and neurological outcomes. We hypothesized that jTBI would increase neuroinflammation and impair development of the OXT system while IN-OXT would mitigate inflammatory responses of the secondary injury cascade, resulting in less neuropathology and better functional recovery.

## Experiment 1

### Methods

All experimental procedures were approved by the Southern Illinois University Institutional Animal Care and Use Committee. Subjects were bred in-house from breeders purchased from Harlan Laboratories (Indianapolis, IN). Thirty animals at post-natal day (PND) 28 were housed in pairs under a 12-h light/dark cycle with access to food and water *ad libitum*. Female subjects were selected based on previous findings indicating greater therapeutic potential of OXT treatment in females (27,31).

### Surgery

Animals received either a bilateral frontal controlled cortical impact (fCCI) or sham surgery and were sacrificed on post-injury days (PIDs) 1, 14, or 30. For fCCI, a midline incision was made, followed by a 5 mm diameter craniotomy centered 2 mm anterior to bregma (+4.5 mm to −0.5 mm) and 0 mm from the midline (+2.5mm to −2.5 mm). An electromagnetically activated piston (Leica, St. Louis, MO) attached to a 4 mm impactor probe was used to strike the cortex at 2.25m/s for 500ms after retracting and lowering the probe by 3 mm. Sham surgery replicated the incision procedure without performing the craniotomy.

### Transcardial Perfusion

Following intraperitoneal injection of 1 mL 50% urethane in water, subjects underwent transcardial perfusion. The entire brain was then removed and post-fixed in 4% paraformaldehyde at 4⁰C for 48 hours, followed by immersion in 30% sucrose solution for cryoprotection before sectioning on a cryostat.

### Histochemistry

Brains were sectioned into 40 μm coronal sections using a Leica CM 1860 cryostat. Four slices (+4.2 mm to +1.7 mm from bregma) were obtained from each animal, followed by an immunohistochemistry protocol (32). Primary antibodies anti-OXTR (1:100; Alomone Labs #AVR-013; Figure 1), anti-OXT (1:10,000, ImmunoStar #20068; Figure 2) and anti-Iba1 (microglia; 1:1000; Wako #019-19741; Figure 3) were used. Slices stained with anti-OXTR were counterstained using cresyl violet (32).

**Figure 1.**
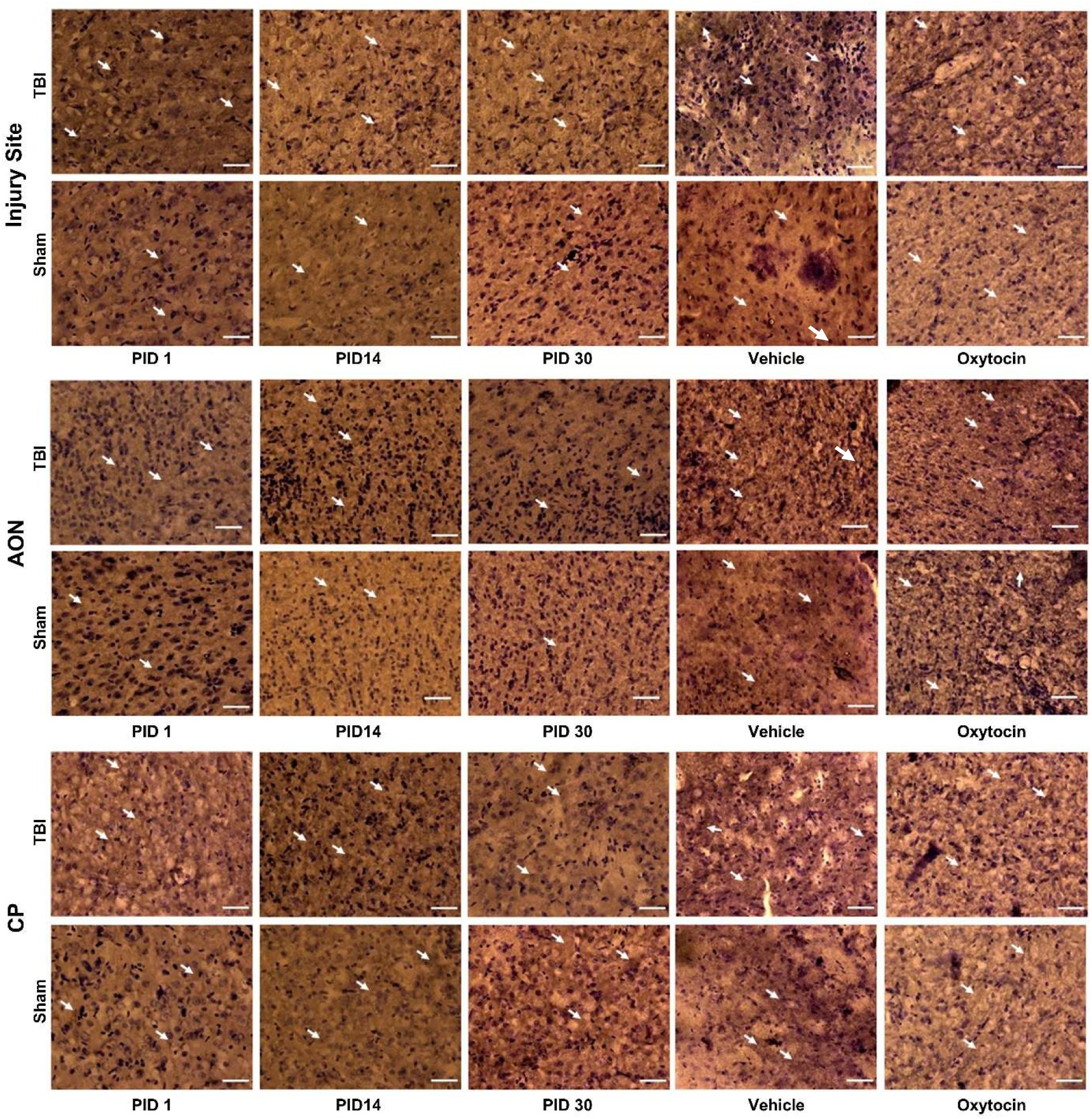
Representative OXTR Images *Note.* Scale bar = 40 μm

**Figure 2.**
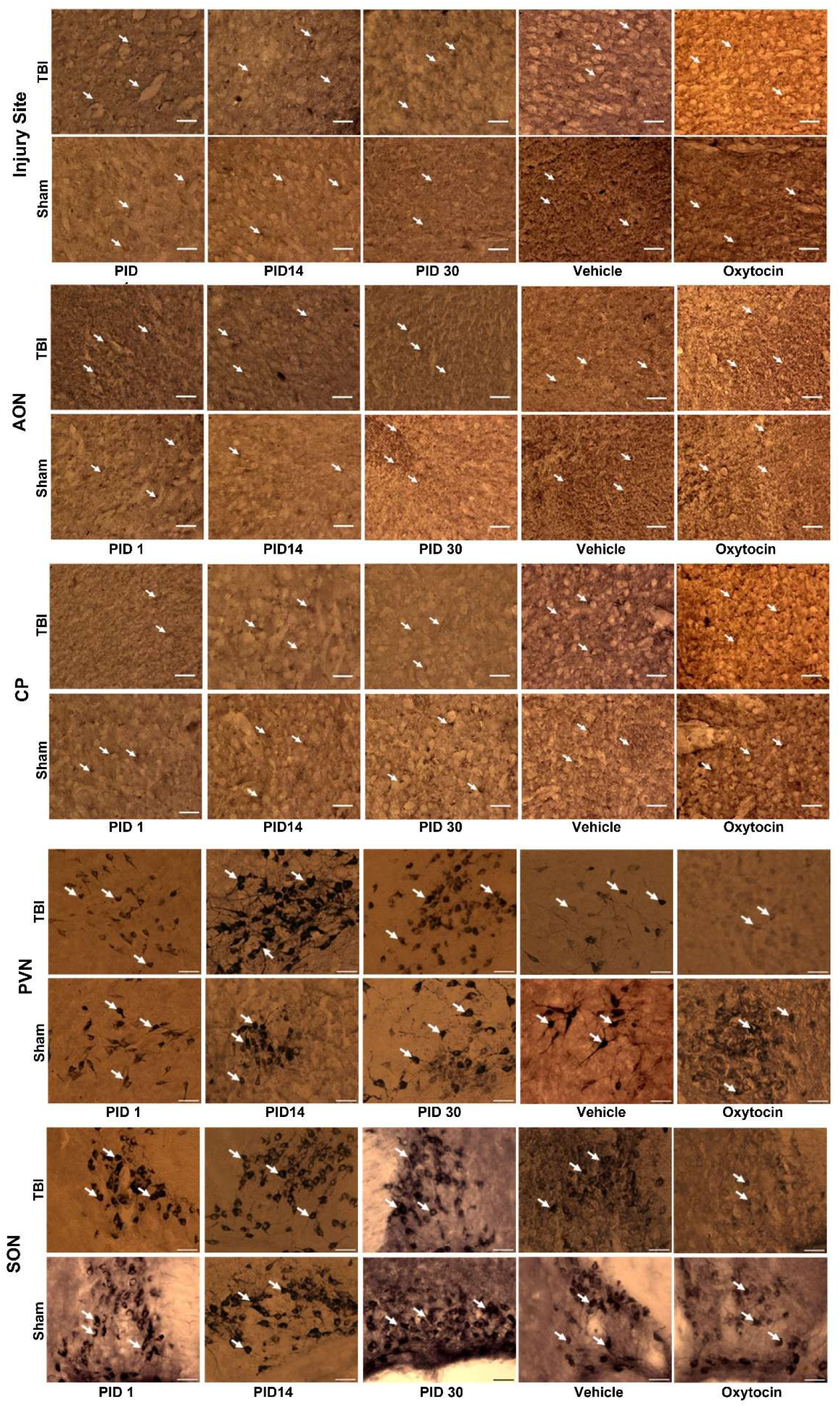
Representative OXT Images *Note.* Scale bar = 40 μm

**Figure 3.**
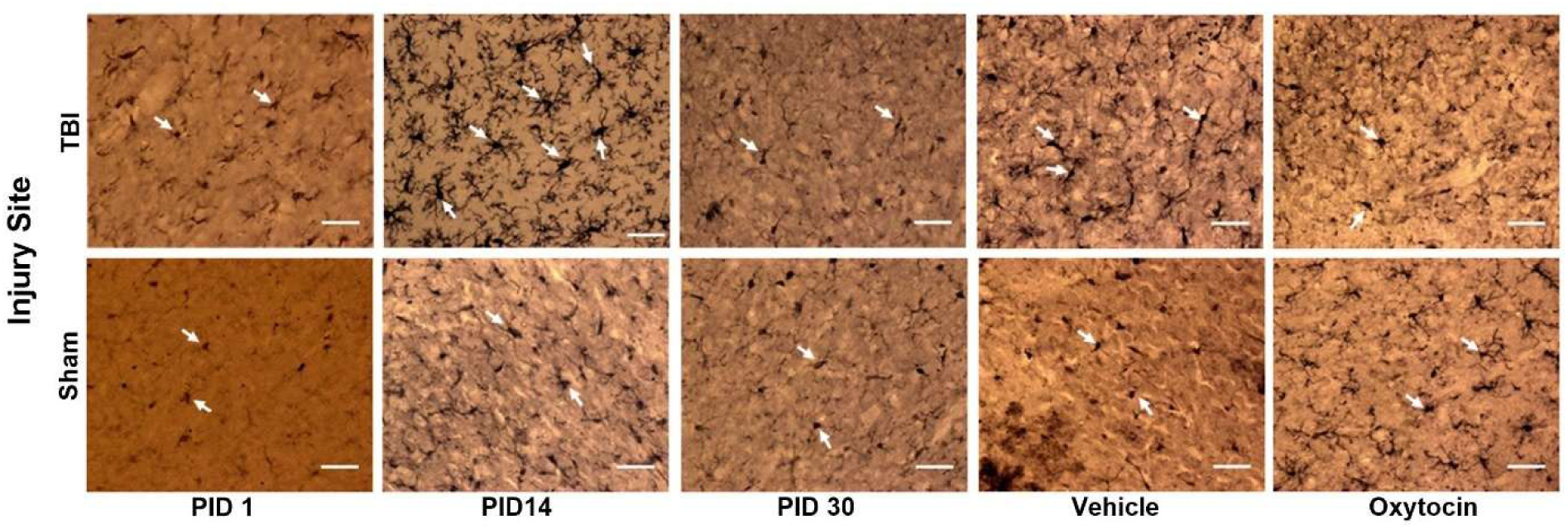
Representative Iba1+ Images. *Note.* Scale bar = 40 μm

For each stain, eight images of the pericontused cortex were taken at the injury site at 40x magnification (60 μm) using an Olympus BX-51 microscope. This region was chosen in order to assess the inflammatory response and subsequent changes in OXT receptor (OXTR) and OXT peptide levels. Additionally, two images of the paraventricular nucleus (PVN) and supraoptic nucleus (SON) of the hypothalamus stained for OXT were taken to quantify TBI-related changes in OXT levels in the brain regions where it is synthesized (23,33–36). While previous research found no effect of injury on OXT mRNA expression in the PVN (25), we will be assessing TBI-related changes to peptide levels. Notably, the SON remains an understudied region, despite being one of the areas where OXT is synthesized and having connections with the forebrain.

Finally, fours images each of the anterior olfactory nucleus (AON) and the caudate putamen (CP) stained for OXT and OXTR were taken to quantify TBI-related changes. These regions were chosen due to their relevance in IN-OXT passage into the brain (37,38) and endogenous OXT production (23,33–36). The images were quantified using ImageJ Fiji (1.54p) where Iba+ cells were manually counted, OXT signal intensity was measured using mean grey values, and OXTR signal intensity was measured using mean grey values that were normalized by the number of cells in the image.

### Statistical Analysis

To assess the microgliosis and the density of OXT and OXTRs, 2 (injury) x 3 (Day: PID 1, 14, and 30) ANOVAs were used to analyze group differences across development. Bonferroni corrected post-hoc tests were then used to identify significant mean differences.

### Experiment 1 Results

All data met the assumptions of normality and homogeneity of variance for the ANOVA tests. Nonsignificant results are included in the supplementary material.

### Microgliosis

TBI significantly impacted microglia [*F*(1, 218) = 1372.677, *p* < 0.001, η^2^ = 0.369] and increased Iba+ cells in TBI animals compared to sham animals (Figure 4; *p*<0.001). This pattern was consistent across development [*F*(2, 218) = 632.509, *p* < 0.01, η^2^ = 0.055] with more Iba+ cells observed in TBI animals (*p*<0.001) that peaked 30 days post-injury (*p*<0.05). Thus, illustrating that TBI progressively increased neuroinflammation at the injury site.

**Figure 4.**
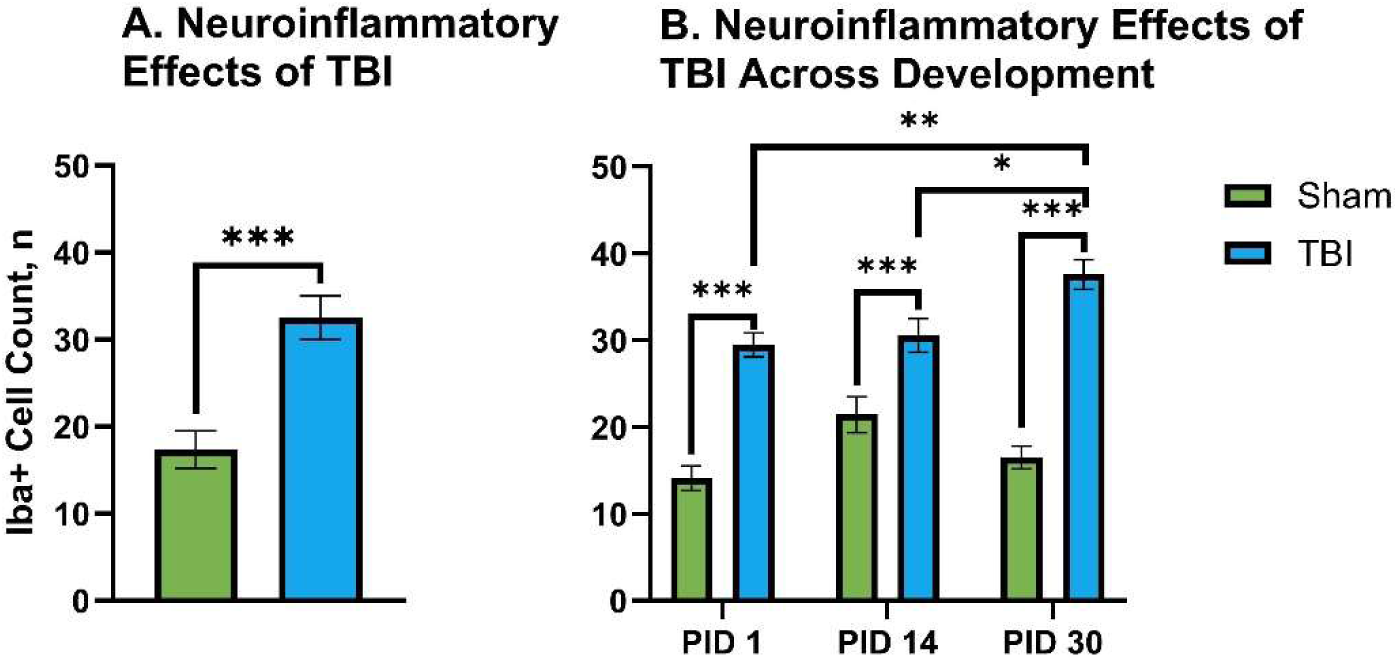
Neuroinflammatory Effects of TBI *Note.* Graphs show group means ± SEM. A) TBI increased inflammation at the injury site, B) TBI animals displayed consistent inflammation across development and it peaked 30 days post-injury. *, p<0.05, **, p<0.01, ***, p<0.001.

### Oxytocin System

At the injury site, TBI did not significantly impact OXTR levels [*F*(1, 218) = 0.564, *p* = 0.453, η^2^ = 0.003] (Figure 5A), but did alter OXTR levels across development [*F*(2, 218) = 5.875, *p* < 0.01, η^2^ = 0.051] (Figure 5B). Specifically sham animals displayed peak OXTR levels in the mPFC on PND 42 (PID 14, *p*<0.001) that subsequently decreased by PND 58 (PID 30, *p*<0.05), thus TBI altered the normal changes in OXTR levels across development. Conversely, TBI did impact OXT levels in this region [*F*(1, 218) = 14.719, *p* < 0.001, η^2^ = 0.063] (Figure 6A), but OXT levels were not affected across development [*F*(2, 218) = 1.435, *p* = 0.240, η^2^ = 0.013] (Figure 6B). Specifically, TBI decreased OXT levels at the injury site (*p*<0.001). Overall, OXTR levels at the mPFC display an inverted U pattern across development and TBI disrupted this pattern, possibly by dampening OXT levels.

**Figure 5.**
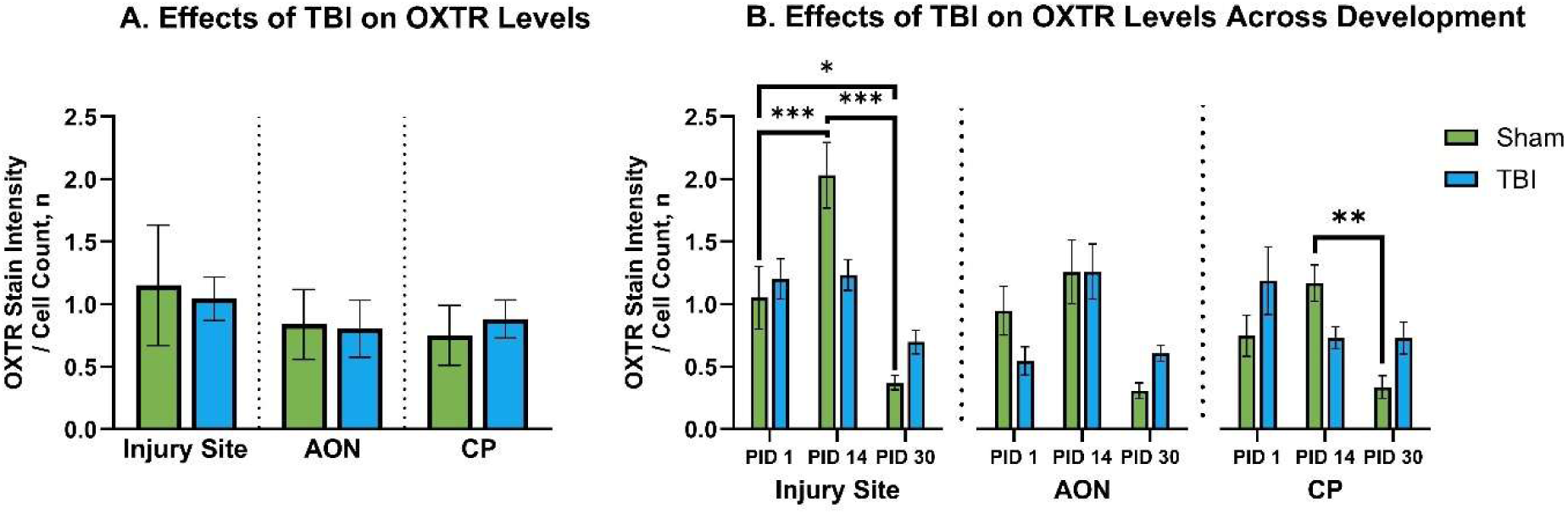
Effects of TBI on OXTR Levels *Note.* Graphs show group means ± SEM. A) TBI did not affect OXTR levels at the injury site, AON or CP, B) Sham animals displayed development differences in OXTR levels, in the mPFC and CP, OXTR levels peaked 14 days following injury. *, *p*<0.05, **, *p*<0.01, ***, *p*<0.001.

**Figure 6.**
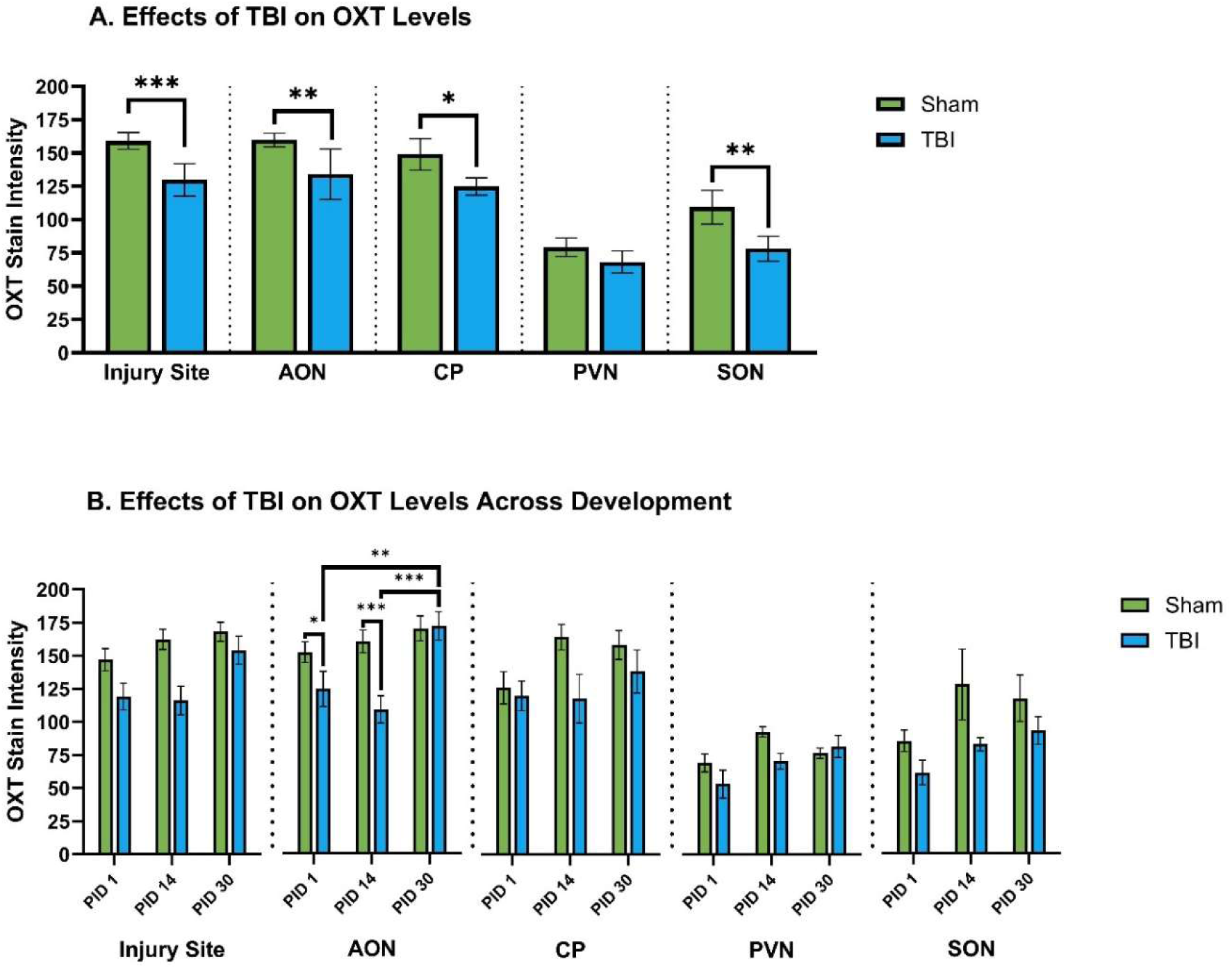
OXT levels across development. Note. Graphs show group means ± SEM. A) TBI decreased OXT levels at the injury site, AON, CP, and SON but not in the PVN, B) TBI decreased OXT levels in the AON on PIDs 1 and 14 but there were no other significant effects of development on OXT levels. *, p<0.05, **, p<0.01, ***, p<0.001.

In the AON, TBI did not significantly impacted OXTR levels [*F*(1, 106) = 0.059, *p* = 0.809, η^2^ = 0.001], nor alter differences across development [*F*(2, 106) = 2.229, *p* = 0.113, η^2^ = 0.04]. However, it did impact OXT levels [*F*(1, 106) = 9.365, *p* < 0.01, η^2^ = 0.081] across development [*F*(2, 106) = 2.229, *p* = 0.113, η^2^ = 0.04] in this region. TBI decreased OXT levels (*p*<0.01) on PIDs 1 (*p*<0.05) and 14 (*p*<0.001) but recovered to sham levels by PID 30 (*p*<0.01). Thus, while OXTR levels were not altered by TBI, it did lower OXT levels for at least two weeks following injury.

Similar to the injury site, TBI did not significantly affect OXTR levels in the CP [*F*(1, 106) = 0.952, *p* = 0.331, η^2^ = 0.009] but did influence the developmental changes in this region [*F*(2, 106) = 4.424, *p* < 0.05, η^2^ = 0.077]. Once again, sham animals displayed increased OXTR levels on PND 42 (PID 14, *p*<0.01) that was not observed in TBI animals. Conversely, OXT levels in the CP mimic those observed at the injury site whereby TBI impacted OXT levels [*F*(1, 106) = 4.710, *p* < 0.05, η^2^ = 0.043] but did not alter the developmental trajectory of OXT levels [*F*(2, 106) = 1.140, *p* = 0.324, η^2^ = 0.021] such that TBI decreased OXT levels (*p*<0.05). Thus, TBI dampens OXT levels, potentially contributing to disruptions to the developmental OXTR pattern.

Finally, TBI did not impact OXT levels [*F*(1, 50) = 3.566, *p* = 0.065, η^2^ = 0.067] or alter the developmental trajectory of OXT levels in the PVN [*F*(2, 50) = 1.956, *p* = 0.152, η^2^ = 0.073]. However, TBI did affect OXT levels in the SON [*F*(1, 50) = 7.505, *p* < 0.01, η^2^ = 0.131] without altering normal developmental changes [*F*(2, 50) = 0.374, *p* = 0.690, η^2^ = 0.015]. Specifically, TBI decreased OXT levels in the SON (*p*<0.01), thus identifying the potential mechanistic origin for OXT deficits observed in the forebrain.

### Experiment 1 Discussion

As hypothesized, TBI increased neuroinflammation across development, persisting for at least 30 days post-injury (28,39). However, despite OXTR being expressed on microglia, TBI did not alter OXTR levels at the injury site, however it did decrease OXT levels throughout the brain, potentially contributing to the unabated microglial response. The observed OXT deficits were likely due to decreased OXT production in the SON but not the PVN. To identify if these deficits could be resolved and inflammation mitigated, we subsequently investigated whether acute IN-OXT administration post-injury could restore depleted OXT levels and promote functional recovery following TBI.

## Experiment 2

### Methods

#### Surgery

For this experiment, 48 animals at PND 28 were utilized and divided into six groups: receiving either fCCI or sham surgery combined with IN-OXT, vehicle (VEH), or no treatment (NoTx). Identical procedures for fCCI and sham surgeries were employed as in Experiment 1.

#### IN-OXT Administration

Animals received IN-OXT, (OXT, Cayman Chemical Company, #11799) diluted in sterile Ringer’s solution (mM: 147.1 Na^+^, 2.25 Ca^2+^, 4 K^+^, 155.6 Cl^-^, pH 7.4) to a concentration of 1 μg/1 μl (20μL) (23,29,36,40–42). VEH animals received an equivalent volume of sterile Ringer’s solution. IN-OXT was administered bilaterally on the rhinarium immediately after scalp closure and before anesthesia recovery, adapting application methods from existing literature (23,41,42).

#### Behavior

The behavior tasks included the modified foot fault task (mFFT), open field (see supplementary material), Morris water maze (MWM), social preference task (see supplementary material), and social dominance task (SDT). For reference Table 1 shows the post-injury days relative to the animal’s age (post-natal day).

**Table 1.**
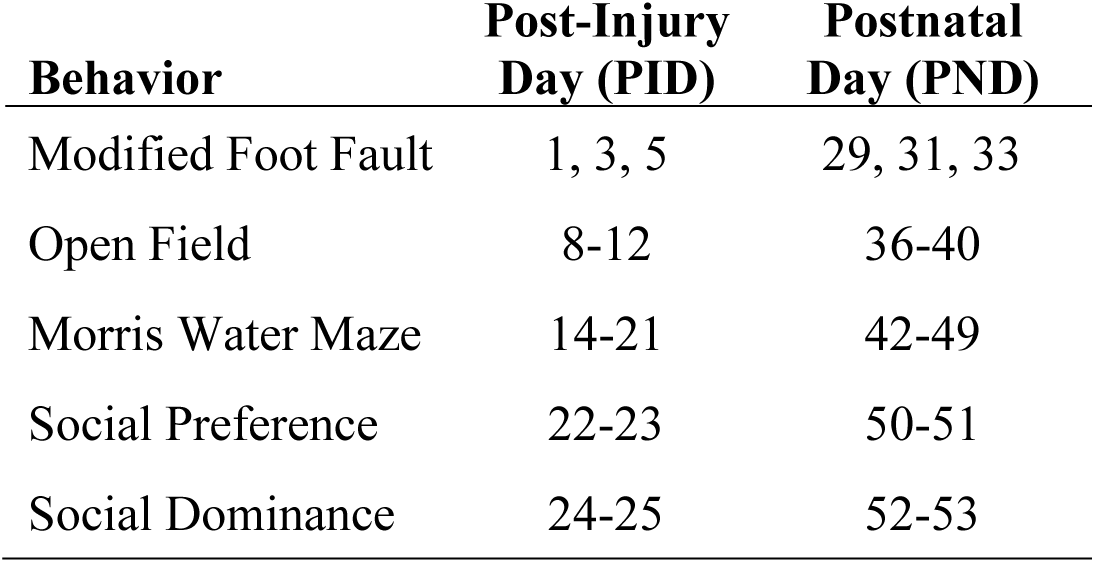
Post-Injury Days Relative to Post-Natal Days.

##### Modified Foot Fault Task

The foot fault task is a standard assessment in TBI research, ensuring motor coordination but without any movement incentives, thus potentially masking deficits (43–47). To address this limitation, our study employed a mFFT, which is a combination of the beam task and foot fault task that was developed in our lab. The beam was 120 cm long and consisted of a 2.5 cm^2^ plastic grid beam attached to a black box.

##### Morris Water Maze

The MWM featured a 200 cm (D) x 74 cm (H) polyethylene pool divided into four quadrants, filled with opaque water. A 13 cm (D) x 72 cm (H) clear plexiglass platform, submerged 2 cm under the water’s surface, was located in the northeast quadrant.

ANY-maze video tracking software recorded animal movements. The MWM comprised three phases: training, probe, and reversal training. These phases followed previously established protocols (32).

Platform search strategy was assessed during training and probe trials, focusing on the progression from inefficient to efficient strategies. Animal coordinates from ANY-maze were analyzed using Pathfinder software (48) to assess search strategies for each trial.

##### Social Dominance Task

The SDT assessed changes in social hierarchy post-TBI and followed a protocol from previous research (49). ‘Winning’ was defined as reaching the entrance barrier slit at the tube’s opposite end, and results were recorded for each target rat. Novel rats were matched in sex and age.

### Histochemistry

The same procedure for transcardial perfusion and immunohistochemistry in experiment 1 was used on animals in experiment 2. Animals were sacrificed on PID 30.

### Statistical Analysis

We collapsed sham groups into a single sham group, additionally we collapsed NoTx TBI and VEH TBI groups into a single NoTx group because we did not find any significant differences between these groups. Analysis involved repeated-measures ANOVAs, MANOVAs and Kruskal-Wallis H Tests. Bonferroni corrected post-hoc tests were employed to identify significant mean differences between groups.

### Experiment 2 Results

All data meet the assumption of normality and homogeneity of variance for the (M)ANOVA tests. Nonsignificant results are included in the supplementary material (i.e., open field and social preference task).

### Modified Foot Fault Task

Treatment status significantly affected foot fault frequency [*F*(2, 45) = 3.512, *p*<0.05, η^2^=0.135] across trial days [*F*(2.729, 61.409) = 3.494, *p*<0.05, η^2^=0.134] (Supplementary Figures). Specifically, TBI animals, regardless of treatment status displayed a higher frequency of foot faults compared to sham animals (*p*<0.05), but these effects were only observed on the final testing day (*p*<0.05). Thus, TBI led to motor coordination deficits and IN-OXT did not attenuate this result.

### Morris Water Maze

#### Training Phase

Treatment status also significantly affected spatial learning [*F*(2, 45) = 25.287, *p*<0.001, η^2^=0.529] across trials days [*F*(6.645, 149.517) = 2.471, *p*<0.05, η^2^=0.099] (Figure 7A). Specifically, NoTx TBI animals exhibited the slowest latency to the platform compared to sham (*p*<0.001) and OXT TBI animals (*p*<0.05), which was consistent across trial days (*p*<0.01). Conversely, TBI animals treated with OXT only displayed a slower latency than sham animals on training days 1 (PID 15; *p*<0.05) and 2 (PID 16; *p*<0.05), and then demonstrated a faster latency than NoTx TBI animals on the final training day (day 5, PID 19; *p*<0.001). Thus, implicating an acute IN-OXT as mildly effective at reversing spatial learning deficits observed following TBI.

**Figure 7.**
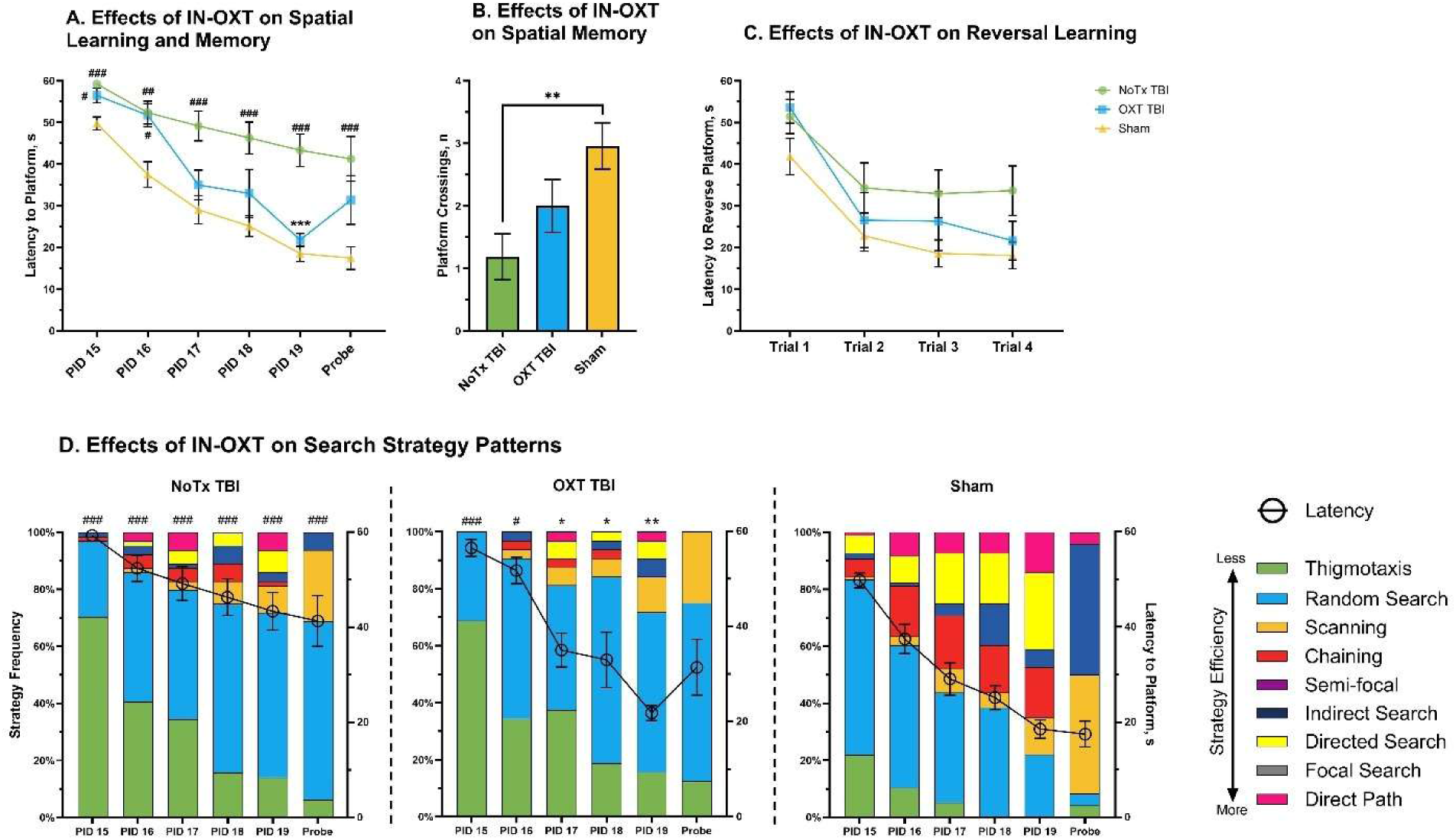
Effects oan IN-OXT on Spatial Learning and Memory in the MWM *Note.* Graphs show group means ± SEM. A) Untreated TBI animals consistently showed impairments in learning and memory compared to sham animals, while OXT TBI animals only showed impairment on PID 15 and 16 and demonstrated no deficits on the last day of training (PID 19), B) NoTx TBI animals displayed spatial memory deficits with fewer platform crossings in the Probe Trial, C) NoTx TBI animals displayed deficits in reversal learning across trials (*p*<0.01) but there were no significant group differences between trials, D) NoTx TBI animals displayed higher levels of inefficient search strategies versus shams on all trial days and the probe trial, OXT TBI animals displayed more inefficient search strategies versus shams on the first two trial days (PID 15, 16) and then displayed more efficient search strategies versus NoTx TBI animals on the remaining trial days (PID 17-19). #, significantly different from sham animals, *, significantly different from NoTx TBI animals, */#, *p*<0.05, **/##, *p*<0.01, ***/###, *p*<0.001

The search strategies of the animals displayed a similar pattern as the latency results, with treatment status significantly impacting search strategy on all training days (PIDs 15 [H(2) = 42.499, *p*<0.001, η^2^ = 0.214], 16 [H(2) = 25.964, *p*<0.001, η^2^ = 0.127], 17 [H(2) = 27.619, *p*<0.001, η^2^ = 0.136], 18 [H(2) = 30.231, *p*<0.001, η^2^ = 0.149], and 19 [H(2) = 37.425, *p*<0.001, η^2^ = 0.187], Figure 7D). Overall, untreated TBI animals consistently displayed less efficient search strategies than sham animals (*p*<0.001). However, OXT treatment mitigated this with OXT TBI animals only displaying less efficient searching than sham animals on training days 1 (PID 15, *p*<0.001) and 2 (PID 16, *p*<0.05), and more effective searching on training days 3 (PID 17, *p*<0.05), 4 (PID 18, *p*<0.05), and 5 (PID 19, *p*<0.01) compared to untreated TBI animals. Thus, acute IN-OXT treatment improves spatial learning via preservation of problem-solving abilities.

#### Probe Trial

During the probe trial, spatial memory was significantly affected by treatment status as measured by the latency to initial platform zone crossing [*F*(1, 45) = 124.396, *p* < 0.001, η^2^ = 0.734] (Figure 7A) and the number of platform zone crossings [*F*(1, 45) = 63.179, *p* < 0.001, η^2^ = 0.584] (Figure 7B). TBI produced significant spatial memory deficits with longer latencies to initial platform zone crossing (*p*<0.001) and fewer platform zone crossings compared to sham animals (*p*<0.01). However, IN-OXT did not mitigate these deficits, despite imparting mild improvements to spatial learning. These results are replicated in search strategy results [H(2) = 16.469, *p* < 0.001, η^2^ = 0.322] (Figure 7D), whereby NoTx TBI animals displayed less effective search strategies compared to sham animals (*p*<0.001). OXT TBI animals did not significantly differ from either group, implying that IN-OXT treatment does not preserve spatial memory following TBI.

#### Reversal Day

After the probe day, animals underwent a test for cognitive flexibility, where the platform was relocated to a new position, and the latency to reach the new platform location was recorded. Treatment status significantly affected reversal learning [*F*(2, 45) = 5.216, *p* < 0.01, η^2^ = 0.188], but did not affect reversal learning across trials [*F*(6, 135) = 0.373, *p* = 0.895, η^2^ = 0.016] (Figure 7C). Specifically, untreated TBI animals displayed slower latencies to the reverse platform location compared to sham animals (*p*<0.01) but IN-OXT treatment did not significantly improve reversal learning abilities.

### Social Dominance Task

Social dominance, as measured by percentage of winning trials, was significantly affected by group [H(2) = 13.131, *p*<0.001, η^2^ = 0.247] (Figure 8). Sham animals consistently displayed fewer winning trials than NoTx TBI (*p*<0.01) and OXT TBI animals (*p*<0.05). Therefore, TBI increased social dominance behaviors, but IN-OXT did not mitigate these effects.

**Figure 8.**
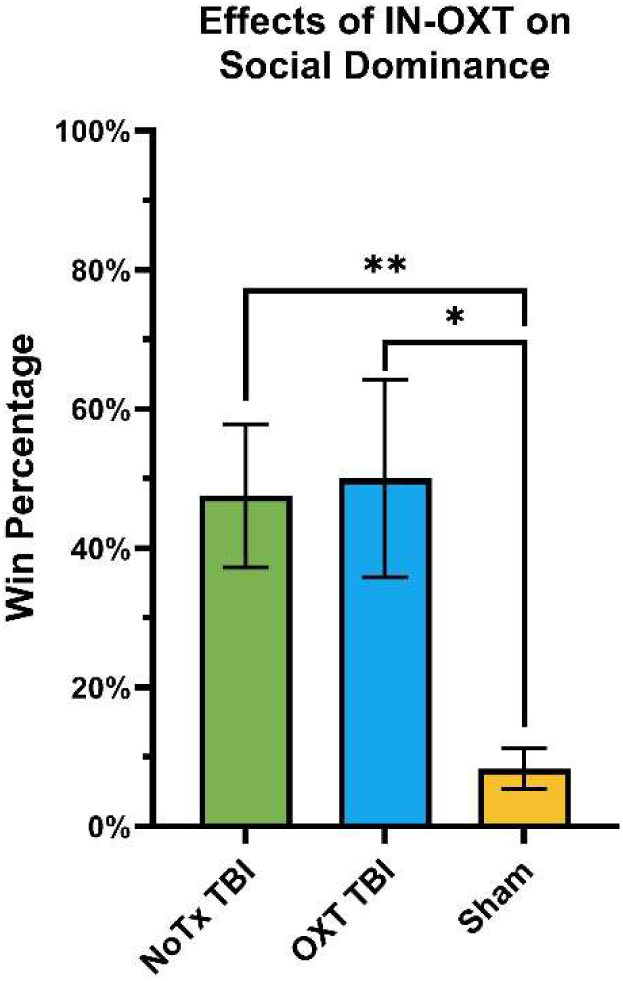
Effects of IN-OXT on Social Dominance *Note.* Graphs show group means ± SEM. TBI animals regardless of treatment status displayed higher levels of social dominance than sham animals. *, *p*<0.05, **, *p*<0.01.

### Microgliosis

To identify the long-term impact of an acute IN-OXT treatment on neuroinflammation, Iba+ cells were counted at the injury site and significant group effects were observed [H(2) = 79.672, *p*<0.001, η^2^ = 0.344] (Figure 9). Sham animals displayed fewer Iba+ cells than NoTx TBI (*p*<0.001) and OXT TBI animals (*p*<0.001). Thus, TBI increased inflammation but IN-OXT did not impart long-term mitigation of the neuroinflammatory response.

**Figure 9.**
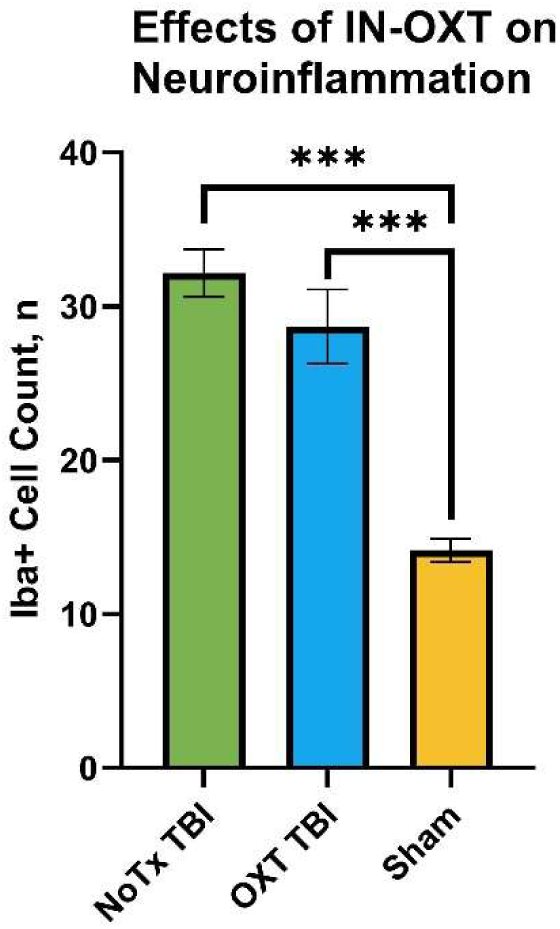
Effects of IN-OXT on Neuroinflammation *Note.* Graphs show group means ± SEM. TBI increased inflammation but IN-OXT did not mitigate this effect. ***, *p*<0.001.

### Oxytocin System

To identify if an acute dose of IN-OXT could rescue the OXT deficits observed in Experiment 1, OXT levels were measured at the injury site and in the AON, CP, PVN, and SON. Additionally, OXTR levels were measured at the injury site, AON, and CP to further quantify the chronic physiological effects of IN-OXT on the OXT system.

At the injury site, treatment status significantly affected OXTR levels [*F*(2, 229) = 4.586, *p* < 0.05, η^2^ = 0.039] (Figure 10A) such that IN-OXT treatment increased OXTR levels relative to sham animals (*p*<0.01). Furthermore, treatment status also significantly affected OXT levels in this region [*F*(2, 229) = 15.304, *p* < 0.001, η^2^ = 0.118] (Figure 10B), with NoTx TBI animals displaying a deficit compared to shams (*p*<0.001). Thus, IN-OXT rescued OXT levels and due to the presence of OXTRs on microglia, potentially contributed to elevated OXTR expression on the microglia in the region.

**Figure 10.**
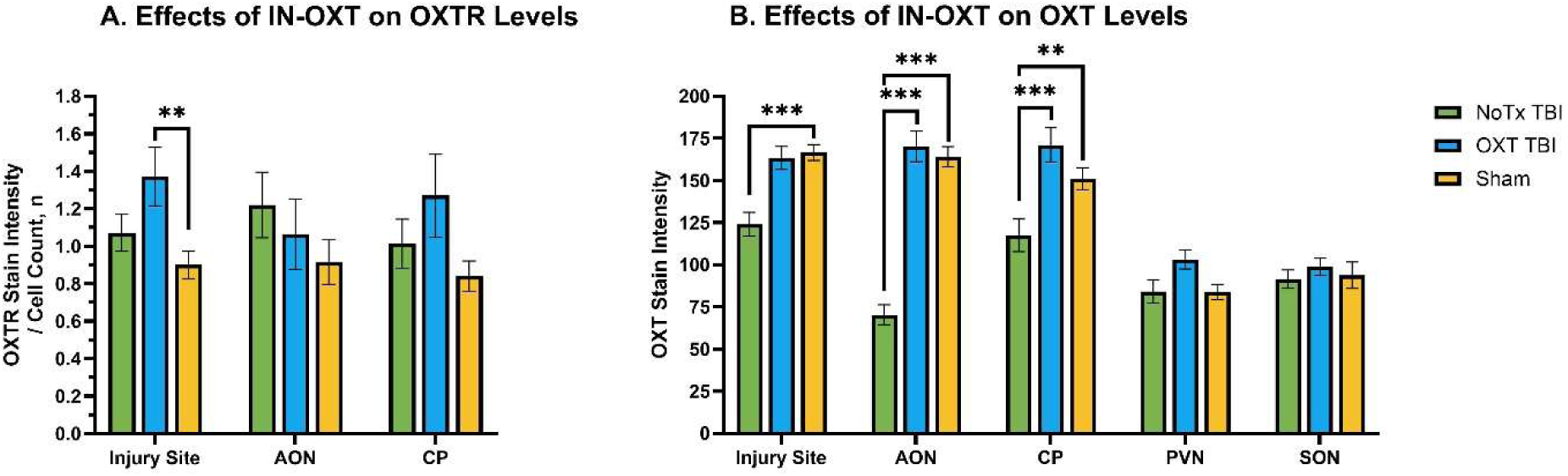
Effects of IN-OXT on the Oxytocin System *Note.* Graphs show group means ± SEM. A) IN-OXT increased OXTR levels at the injury site, B) TBI decreased OXT levels at the injury site and in the AON and CP but IN-OXT treatment mitigated this effect, but there was no effect of TBI or IN-OXT on OXT levels in the PVN or SON. **, *p*<0.01, ***, *p*<0.001.

In the AON, while IN-OXT treatment did not impact OXTR levels [*F*(2, 113) = 1.156, *p* =0.318, η^2^ = 0.02], it did affect OXT levels [*F*(2, 113) = 7.847, *p* < 0.001, η^2^ = 0.122]. Similar to the injury site, NoTx TBI animals displayed decreased levels of OXT compared to (*p*<0.001) and OXT TBI animals (*p*<0.001). Thus, IN-OXT rescued the OXT deficits in the AON caused by TBI.

As in the AON, OXTR levels in the CP was not affected by treatment [*F*(2, 113) = 2.547, *p* = 0.083, η^2^ = 0.043] but OXT levels were [*F*(2, 113) = 8.138, *p* < 0.001, η^2^ = 0.126]. Once again, the NoTx TBI group displayed deficits compared to sham (*p*<0.01) and OXT TBI (*p*<0.001) animals. Therefore, IN-OXT proves to be a useful treatment for preventing OXT deficits.

Finally, while IN-OXT treatment displayed chronic restoration of OXT levels in the forebrain, no such effects were observed in the PVN [*F*(2, 55) = 2.427, *p* = 0.098, η^2^ = 0.081] or the SON [*F*(2, 55) = 0.142, *p* = 0.868, η^2^ = 0.005]. Thus, while TBI reduces OXT levels in the forebrain, it does not impact the synthesis of OXT, suggesting that other mechanisms are involved in the recovery effects of IN-OXT observed in the forebrain.

### Experiment 2 Discussion

Animals with TBI exhibited gross motor deficits, consistent with prior studies (32,44,45), but acute IN-OXT treatment failed to enhance performance. However, IN-OXT did yield modest improvements in spatial learning, as seen in previous research (50), which is remarkable given the large time span between dose delivery and task performance. Unfortunately, these benefits did not extend to spatial memory or reversal learning. Furthermore, while TBI heightened social dominance, IN-OXT did not mitigate this effect. It is possible that administration of IN-OXT is required immediately preceding task performance to observe effects. Indeed, previous research has found that for social tasks specifically, OXT needs to be administered prior to the task in order to observe improvements in social functioning (51,52, reviewed in 53). Overall, IN-OXT imparted only mild enhancements in spatial learning and failed to improve functional recovery of other behaviors.

Histologically, IN-OXT treatment did not alleviate neuroinflammation, despite elevating OXTRs and rectifying OXT deficits at the injury site. Thus, acute IN-OXT proved ineffective on neuroinflammation post-TBI. While IN-OXT did not affect OXTR levels in the AON or CP, it did restore OXT deficits induced by TBI in both regions. Additionally, consistent with Experiment 1 and previous findings (25), TBI had no impact on OXT levels in the PVN. Surprisingly, there were no injury or treatment effects on OXT levels in the SON, diverging from Experiment 1. This suggests that IN-OXT treatment resolves OXT deficits within a month post-injury, returning OXT levels in other brain regions to normal. Further investigation is needed on the acute-phase effect of IN-OXT treatment in the SON.

## Discussion

The study revealed that juvenile TBI induces behavioral deficits (motor, learning, memory, social dominance) linked to increased inflammation, consistent with prior research (46,47,54–57). A novel finding was the elevation of social dominance in young female animals following a focal CCI injury to the mPFC, contrasting with deficits observed in adulthood (58). While frontal injuries are known to increase aggression in males due to deficits in impulse control and executive function (59,60), the acute dosing regimen of IN-OXT was not effective at mitigating these effects. However, acute OXT treatment post-TBI did led to mild enhancements in spatial learning, aligning with previous findings (50), but failed to improve spatial memory recall in subsequent probe trials. More research needs to be done to identify the correct dosing regimen to observe behavioral effects but the promising long-term effects on spatial memory suggest that alterations in dosing schedules of IN-OXT may impart behavioral sparing of spatial learning.

This study also found novel developmental deficits in the OXT system due to TBI. We observed that TBI altered the normal developmental pattern of OXTRs in the mPFC and CP. While the deficits in the mPFC may be due to it being the injury site, the altered developmental trajectory in the CP indicates that TBI alters normal developmental processes of the OXT system in brain regions adjacent to the injury site. Furthermore, TBI caused chronic deficits in OXT levels in the forebrain, likely due to deficits observed in the SON. While both the PVN and SON synthesize OXT and project to forebrain regions (35,61), the PVN remains the primary focus in OXT research despite demonstrating no observable effects of TBI (52). However, our results suggest that OXT deficits in the forebrain may be due to decreases in the synthesis of the peptide in the SON. These deficits may only last until 30 days post-injury as Experiment 2 did not observe any effects of TBI on OXT levels in this region. Nonetheless, more research is necessary to confirm this mechanistic effect as SON results in Experiment 1 were not consistent in Experiment 2.

Despite these inconsistencies, we observed significant histological effects of IN-OXT following TBI. While IN-OXT did not alleviate inflammation as expected, it did increase OXTR levels at the injury site. While research that colabels microglia with OXTR needs to be done to identify if this increase is related to the increased inflammatory response, these results suggest that IN-OXT increases OXTR expression at the injury site, thereby restoring OXT levels to normal. Thus, follow-up studies are needed to better identify the role IN-OXT plays in inflammation as an acute dose may not be enough to impart significant long-term mitigation.

Despite this, IN-OXT appears to restore OXT levels in the forebrain, that may be due to mechanisms of the SON. However, since OXT levels in this region did not differ between untreated and OXT-treated TBI animals, this suggests that OXT synthesis was not altered by either TBI or IN-OXT. Thus, more research is necessary to identify the mechanism by which forebrain OXT levels are chronically altered.

Overall, OXT-treated TBI animals exhibited mild spatial learning improvements, possibly attributed to hippocampal effects (50). However, IN-OXT failed to ameliorate social dominance changes, despite its effects on OXT and OXTR levels in the frontal lobe. Further research into additional brain regions is necessary to elucidate these deficits. Lastly, the SON, not the PVN, seems responsible for OXT system deficits in the frontal lobe. However, since IN-OXT resolved OXT deficits without attenuating inflammation, exploration of chronic dosing is recommended.

