## Supplementary Material for "Acute Administration of Oxytocin in the Functional Recovery of Neurocognitive and Social Deficits Following Juvenile Frontal Traumatic Brain Injury"

### Methods

#### *Open Field Task*

The open field (OF) task consists of a 50-cm x 50-cm box with a camera tracking the animal from above using ANY-maze software. Testing consisted of one trial per day for five days starting on PID 8. ANY-maze software measured the rat's location in the center and at the edges, as well as the time spent immobile. A location ratio was calculated using location information (outer edge time – center time / total time).

#### *Social Preference Task*

The social preference task (SPT) was performed in a clear plexiglass box (144 cm x 34 cm x 26 cm) with three chambers in a room with minimal lighting and the task began on PID 22. The three chambers consisted of a central compartment where the rat was initially placed as well as a left and right compartment. The procedure followed a protocol outlined in previous research (1). The amount of time the rat interacted with the novel rat and the novel object was recorded with a video camera. Interactions are measured by the amount of time that the test rat spent at the barrier by the novel rat (nose-touching, sniffing, climbing).

#### *Lesion Verification*

Sections for lesion verification were stained using cresyl violet and captured with an Olympus BX-51 microscope and Qimaging Retiga R6 camera. The areas of the lesioned tissue were measured by hand using ImageJ. Total cortical area was calculated by adding the two hemispheres together. The Calvalieri method was then used to calculate the volume (2).

### Results

#### *Open Field*

For the open field, there were no significant group effects for time spent immobile [$F$(5.511, 124.003) = 0.859, $p$ = 0.519, $\eta^2$=0.037], or location ratio [$F$(8, 180) = 0.815, $p$ = 0.59, $\eta^2$=0.035] (Figure S2). Thus, neither TBI nor IN-OXT had an effect on anxiety-like behavior.

***Social Preference Task***

For the social preference task, there were no significant group effects for the number of interactions with the novel object [$F$(2, 45) = 0.369, $p$=0.693, $\eta^2$=0.016], nor novel rat [$F$(2, 45) = 2.466, $p$=0.096, $\eta^2$=0.099] (Figure S3). There were also no significant group effects for the time spent interacting with the novel object [$F$(2, 45) = 1.024, $p$=0.367, $\eta^2$=0.044], nor novel rat [$F$(2, 45) = 0.559, $p$=0.576, $\eta^2$=0.024]. Thus, neither TBI nor IN-OXT affected social preferences.

***Lesion Analysis***

When measuring the progression of the lesion across development (Figure S4), there was a significant effect of injury [$F$(1, 22) = 44.854, $p$<0.001, $\eta^2$=0.671] (Figure S5A), but not injury*day [$F$(2, 22) = 0.651, $p$=0.531, $\eta^2$=0.056] (Figure S5B). Sham animals displayed more cortical volume than TBI animals ($p$<0.001). Next, we measured whether IN-OXT treatment mitigated lesion size and found a significant effect of group [H(2) = 14.417, $p$<0.001, $\eta^2$=0.478] (Figure S5C). Sham animals displayed more cortical volume than NoTx TBI animals ($p$<0.01) and OXT TBI animals ($p$<0.05). Thus, TBI resulted in cortical tissue loss but IN-OXT treatment did not mitigate this.

1. Chou A, Morganti JM, Rosi S. Frontal Lobe Contusion in Mice Chronically Impairs Prefrontal-Dependent Behavior. PLoS One. 2016/03/11 ed. 2016;11(3):e0151418.

2. Coggeshall RE. A consideration of neural counting methods. Trends Neurosci. 1992/01/01 ed. 1992 Jan;15(1):9–13.

3. Etehadi Moghadam S, Azami Tameh A, Vahidinia Z, Atlasi MA, Hassani Bafrani H, Naderian H. Neuroprotective Effects of Oxytocin Hormone after an Experimental Stroke Model and the Possible Role of Calpain-1. J Stroke Cerebrovasc Dis. 2017/12/19 ed. 2018 Mar;27(3):724–32.

### Supplementary Figures

**Figure S1. Effects of TBI and IN-OXT on Gross Motor Skills.**

*Note.* Graphs show group means ± SEM. A) TBI decreased gross motor skills and IN-OXT did not mitigate this effect, B) TBI animals only showed gross motor deficits on PID 5 and IN-OXT did not mitigate this effect. *, $p<0.05$.

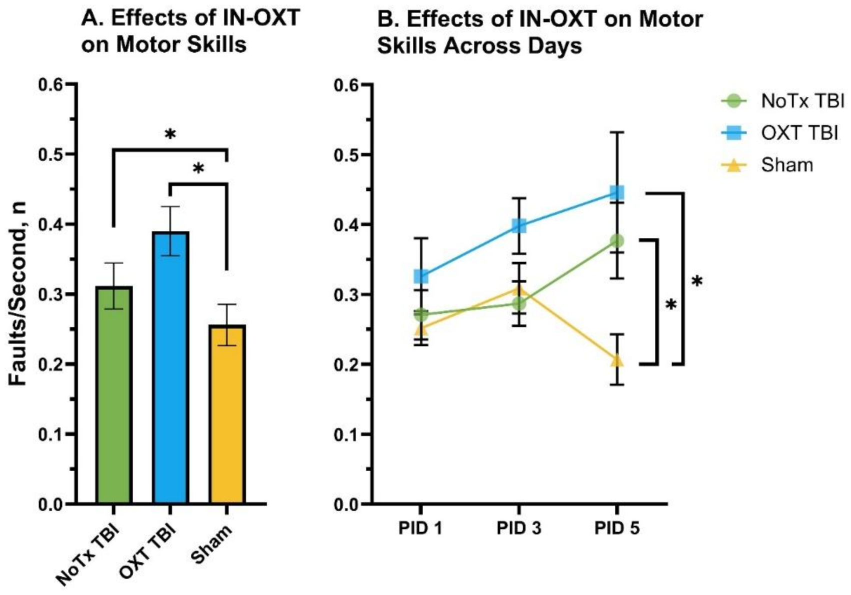


**Figure S2. Effects of IN-OXT on Anxiety-Like Behavior in the Open Field**

*Note.* Graphs show group means ± SEM. There were no significant effects of TBI or IN-OXT on anxiety-like behavior as demonstrated by A) time spent immobile and B) time spent in the outer edge versus the center of the open field.

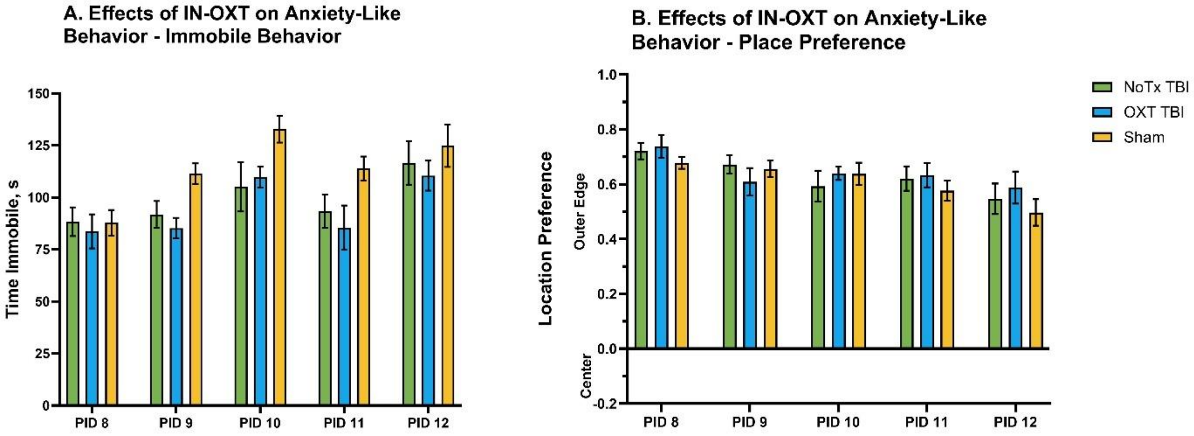


**Figure S3. Effects of IN-OXT on Social Preference**

*Note.* Graphs show group means ± SEM. There were no significant effects of TBI or IN-OXT on social preference behavior as demonstrated by A) number of interactions with either the novel rat or object, and B) time spent interacting with either the novel rat or object.

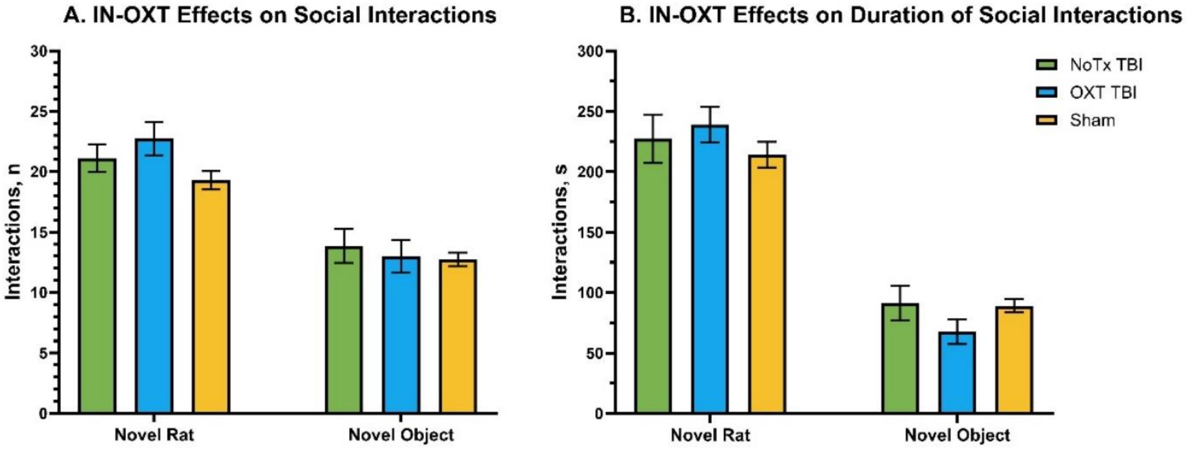


**Figure S4. Representative Cresyl Violet Images**

*Note.* Scale bar = 1 mm

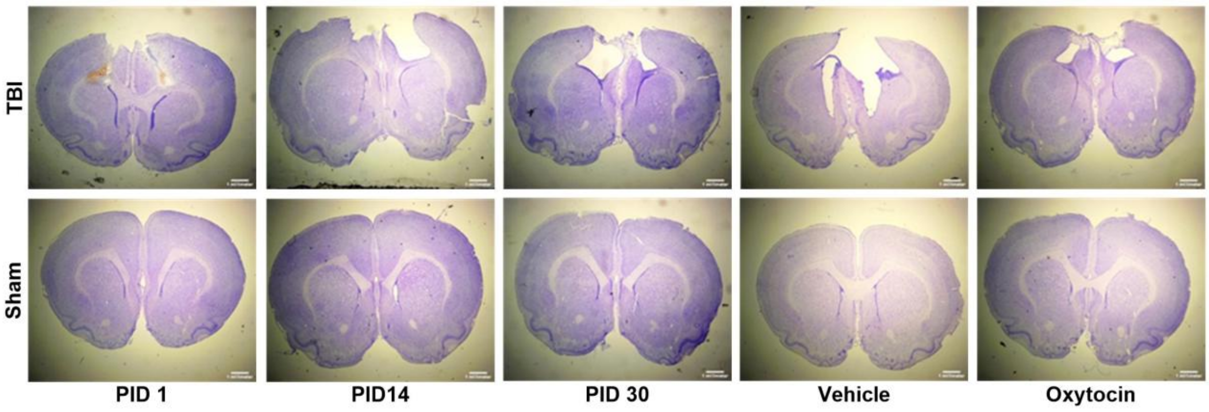


**Figure S5. Effects of TBI and IN-OXT on Cortical Volume**

*Note*. Graphs show group means ± SEM. A) TBI significantly decreased cortical volume but B) there were no differences across development, C) IN-OXT did not mitigate tissue loss. *, $p<0.05$; **, $p<0.01$, ***, $p<0.001$.

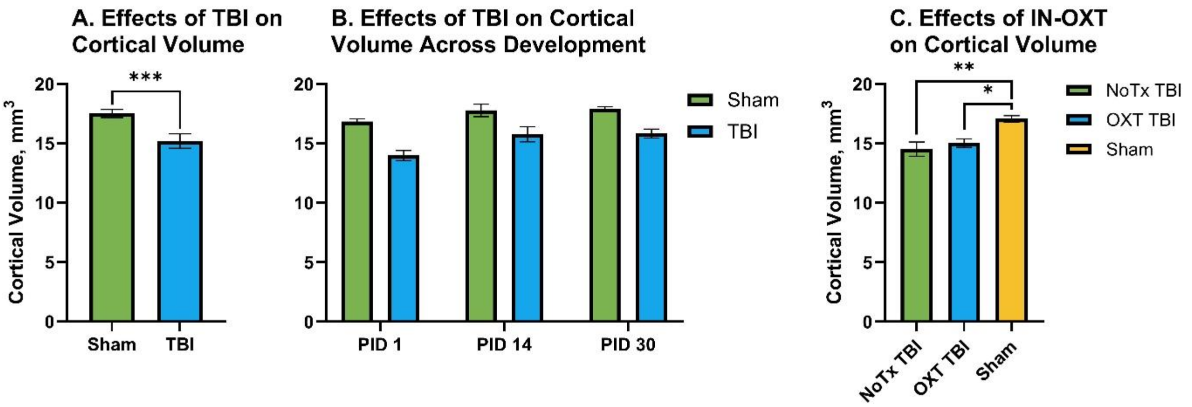
